# Lessons from relatives: C4 photosynthesis enhances CO_2_ assimilation during the low-light phase of fluctuations

**DOI:** 10.1101/2023.04.03.535443

**Authors:** Lucίa Arce Cubas, Cristina Rodrigues Gabriel Sales, Richard L. Vath, Emmanuel L. Bernardo, Angela C. Burnett, Johannes Kromdijk

**Affiliations:** University of Cambridge, Department of Plant Sciences, Downing Street, CB2 3EA Cambridge, United Kingdom; Institute of Crop Science, College of Agriculture and Food Science, University of the Philippines Los Baños, College, Laguna 4031, Philippines; Carl R Woese Institute for Genomic Biology, University of Illinois at Urbana-Champaign, 61801 Urbana, Illinois, USA

## Abstract

Despite the global importance of species with C_4_ photosynthesis, there is a lack of consensus regarding C_4_ performance under fluctuating light. Contrasting hypotheses and experimental evidence suggest that C_4_ photosynthesis is either less, or more efficient in fixing carbon under fluctuating light than the ancestral C_3_ form. Two main issues were identified that may underly the lack of consensus: neglect of evolutionary distance between selected C_3_ and C_4_ species and use of contrasting fluctuating light treatments. To circumvent these issues, we compared photosynthetic responses to fluctuating light across three independent phylogenetically controlled comparisons between C_3_ and C_4_ species from *Alloteropsis*, *Flaveria*, and *Cleome* genera under 21% and 2% O_2_. Leaves were subjected to repetitive stepwise changes in light intensity (800 and 100 µmol m^-2^ s^-1^ PFD) with three contrasting durations: 6, 30 and 300 seconds. These experiments reconcile the opposing results found across previous studies showing that 1) stimulation of CO_2_ assimilation in C_4_ species during the low light phase was both stronger and more sustained than in C_3_ species; 2) CO_2_ assimilation patterns during the high light phase were genus-specific rather than impacted by photosynthetic pathway; and 3) the duration of each light step in the fluctuation regime can strongly influence experimental outcomes.

**One sentence significance statement:** Comparing photosynthesis in three pairs of closely related C_3_ and C_4_ species across three fluctuating light regimes showed that C_4_ photosynthesis has a systematic advantage under the low light phase not related to suppression of photorespiration, while the comparative efficiency under the high light phase was not determined by photosynthetic pathway.

## Introduction

Plants exhibiting C_4_ photosynthesis are mostly found in warm, high light environments (Sage, 2000). Although these environments lead to high light intensity at the top of plant canopies, the light conditions experienced by leaves within the canopy can be extremely dynamic. Indeed, sun angle and cloud cover can alter light intensity by orders of magnitude on a second to minute scale, and shading by higher leaves can further modify the temporal fluctuations experienced by individual leaves. Several C_4_ species form dense canopies with extensive self-shading where sunflecks can provide up to 90% of the energy for photochemistry (Pearcy, 1990, Slattery et al., 2018, Tang et al., 1988, Way and Pearcy, 2012, Zhu et al., 2004). Since photosynthetic responses to changes in light are not instantaneous, fluctuating light has been identified as an area of improvement for crop productivity (Acevedo-Siaca et al., 2020, Kaiser et al., 2018, Kromdijk et al., 2016, Lawson et al., 2012, Long et al., 2006, Pearcy, 1990, Taylor and Long, 2017). Although recent studies have begun to characterise the C_4_ response (Kubásek et al., 2013, Lee et al., 2022, Li et al., 2021, Pignon et al., 2021), most of our understanding of the limitations of photosynthesis under fluctuating light still comes from C_3_ species (Kaiser et al., 2018, Pearcy, 1990, Pearcy et al., 1997). Despite the undeniable global importance of C_4_ crops, with maize and sugarcane alone accounting for over 30% of global agricultural production (FAO, 2020), the impact of the CO_2_ concentrating C_4_ acid shuttle on photosynthetic performance in dynamic light remains understudied (Sales et al., 2021, Slattery et al., 2018) C_4_ photosynthesis is a remarkably ubiquitous adaptation that has evolved independently at least 66 times in angiosperms (Kellogg, 2013) and typically leads to faster photosynthetic rates, higher yields, and greater water use efficiency than the ancestral C_3_ pathway (Kiniry et al., 1989, Sage, 2004). Most C_4_ species operate their carbon concentrating mechanism (CCM) by compartmentalising initial carbon fixation and assimilation between the morphologically distinct mesophyll (M) and bundle sheath (BS) cells, arranged concentrically around the leaf vasculature in ‘Kranz’ anatomy. In the cytosol of M cells, CO_2_ is rapidly converted to bicarbonate by carbonic anhydrase and fixed by phosphoenolpyruvate carboxylase (PEPC) into a 4-carbon oxaloacetate molecule that is further reduced into more stable metabolites malate (Mal) or aspartate (Asp) for transport into the BS. The 4-carbon molecules are decarboxylated in the BS to concentrate CO_2_ around the carbon-fixing enzyme ribulose 1,5-biphosphate carboxylase/oxygenase (Rubisco) and thus enhance photosynthesis by suppressing Rubisco’s alternative oxygenation reaction and resulting photorespiration, which consumes energy and reducing equivalents and re-releases CO_2_ (Leegood, 2002). Reduced carbon in the form of alanine (Ala) or pyruvate is then transported back to the M cells, where phosphoenolpyruvate (PEP) is regenerated at the cost of ATP, imposing an additional cost to C_4_ metabolism. The specific transport metabolites and enzymes of the C_4_ pathway vary, and whilst species were traditionally classified based on predominant decarboxylases nicotinamide adenine dinucleotide-malic enzyme (NAD-ME), nicotinamide adenine dinucleotide phosphate-malic enzyme (NADP-ME), and phosphoenolpyruvate carboxykinase (PEPCK) (Hatch et al., 1975), there is a growing consensus that different decarboxylating enzymes often operate in conjunction, with PEPCK likely acting predominantly as a supplementary pathway (Calsa and Figueira, 2007, Furbank, 2011, Pick et al., 2013, Sales et al., 2018). Crucially, intercellular transport of C_4_ intermediates is driven by diffusion, making the establishment of high metabolic gradients a requirement for the operation of the CCM (Arrivault et al., 2017, Leegood and Furbank, 1984, Lilley et al., 1977, Stitt et al., 1985), although model simulations suggest mixed C_4_ pathways could be less reliant on large metabolite pools (Wang et al., 2012).

Studies conducted on C_3_ species show that the response of CO_2_ assimilation to fluctuating light is restricted by several factors: slow stomatal opening reduces CO_2_ diffusive transfer into the leaf and slow stomatal closing decreases water use efficiency (McAusland et al., 2016), Rubisco activation and the regeneration of ribulose-1,5-biphosphate (RuBP) delay C_3_ cycle activity (Mott and Woodrow, 2000, Pearcy and Seemann, 1990, Sassenrath-Cole and Pearcy, 1992), and the speed of up- and down-regulation of photoprotection lowers light use efficiency (Niu et al., 2022, Zhu et al., 2004). Although said limitations exist irrespective of photosynthetic pathway, C_4_ species have the additional challenge of coordinating the C_3_ and C_4_ cycles (Kromdijk et al., 2014), and perhaps due to this additional complexity, the specific C_4_ response to fluctuating light is not yet fully understood (Kaiser et al., 2018, Slattery et al., 2018). Two apparently contradictory hypotheses can be found in the literature – where C_4_ photosynthesis is suggested to be either less (Kubásek et al., 2013, Li et al. 2021), or more efficient (Stitt and Zhu, 2014) under fluctuating light than the ancestral C_3_ form.

The first hypothesis suggests that C_4_ species are more negatively impacted by sudden changes in light intensity due to the C_3_ and C_4_ cycles temporarily operating asynchronously (Sage and McKown, 2006). Fluctuations in light could disrupt the build-up of metabolic gradients necessary for the effective operation of the CCM, leading to impaired suppression of photorespiration and reduced photosynthetic efficiency (Kromdijk et al., 2010, Slattery et al., 2018). Alternatively, if the CCM is faster to activate during light induction than the C_3_ cycle there could be transient over-pumping of CO_2_ and an increase in BS leakiness – where CO_2_ diffuses out of BS cells back into M cells, raising the energetic cost of carbon fixation due to the futile cycling of PEP. Greater BS leakiness during induction relative to steady state has been previously observed in maize and sorghum (Wang et al., 2022). Lags in CCM deactivation during transitions to lower light could also increase leakiness and reduce quantum yields if Mal/Asp accumulated in the BS is decarboxylated despite insufficient C_3_ cycle activity. In further support for the “negative effect” hypothesis, previous studies have found assimilation rates under fluctuating light relative to steady state to be almost four times lower in C_4_ compared to C_3_ species due to slower photosynthetic induction (Li et al., 2021), as well as a more pronounced reduction of biomass in C_4_ than C_3_ plants grown under fluctuating compared to steady light conditions (Kubásek et al., 2013).

The second hypothesis instead posits that C_4_ species are better able to buffer against sudden changes in light intensity because the large metabolite pools required to drive the CCM can store and release ATP and reducing equivalents (Stitt and Zhu, 2014). The reversible reactions linking the exchange of 3-phosphoglyceric acid (3-PGA) and triose phosphates (TP) between M and BS cells could provide or consume ATP and NADPH to support the C_3_ cycle (Leegood and von Caemmerer, 1989); and mixed C_4_ pathways could transiently enhance Mal over Asp decarboxylation to temporarily increase transport of redox equivalents into the BS (Wang et al., 2014a). In favour of the “positive effect” hypothesis, some of the highest post-illumination CO_2_ fixation rates have been found in C_4_ species (Laisk and Edwards, 1997), and a recent study on grasses recorded higher rates of carbon assimilation in C_4_ over C_3_ species under fluctuating light due to slower decreases in photosynthetic capacity during high-to-low light transitions (Lee et al., 2022).

Although seemingly opposing, there are indications that both hypotheses may coexist. Features of C_4_ biochemistry could have mixed effects – the need to establish large metabolite pools could slow photosynthetic induction but enable higher rates of assimilation upon transitioning to a lower light intensity. Slattery et al. (2018) estimated the buffering capacity of C_4_ photosynthesis to be limited to 10-15 seconds based on maize metabolite pool sampling (Arrivault et al., 2017) and suggested that the specific C_4_ response could thus depend on the length of light fluctuations. A time-sensitive response could account for the different responses observed between sunflecks (Laisk and Edwards, 1997) and longer fluctuations (Kubásek et al., 2013, Li et al., 2021). However, the different light treatments and species used across studies makes it difficult to draw clear conclusions. This is further complicated by C_4_ subtype-specific responses like the post-illumination CO_2_ burst often observed in NADP-ME species (Krall and Pearcy, 1993, Lee et al., 2022), as well as lack of consideration of phylogenetic distance, which can strongly confound comparisons between photosynthetic pathways (Taylor et al., 2010), leading to the inappropriate association of species-specific phenomena with the presence or absence of the C_4_ pathway.

In this study, we compared the photosynthetic response to fluctuating light in relation to steady state across three phylogenetically linked pairs of C_3_ and C_4_ species from *Alloteropsis* (C_3_ *Alloteropsis semialata GMT* & C_4_ *Alloteropsis semialata MDG*), *Flaveria* (C_3_ *Flaveria cronquistii* & C_4_ *Flaveria bidentis*), and *Cleome* (C_3_ *Tarenaya hassleriana* & C_4_ *Gynandropsis gynandra*) genera. Together, these three pairs contain both monocots and dicots, and all three C_4_ decarboxylase subgroups. To compare gas exchange responses across the three pairs, fluctuating light treatments consisting of repetitive stepwise changes in light intensity from 800 to 100 µmol m^-2^ s^-1^ photon flux density (PFD) were used, with three different light step durations: 6, 30, and 300 seconds. To evaluate the impact of photorespiration on the responses to fluctuating light, experiments were conducted under both 2% and 21% oxygen concentration. We hypothesised that 1) after a transition to low light, C_4_ species will be better able to sustain photosynthetic rates than C_3_ species, even when photorespiration is suppressed, 2) C_4_ species will incur a higher penalty during transitions to high light; and 3) the impact of both effects will be inversely correlated to fluctuation length.

## Results

### Steady state responses of CO_2_ assimilation in paired C_3_ and C_4_ species are consistent with well-established differences between photosynthetic pathways

To provide a baseline for comparing CO_2_ assimilation rates between three pairs of closely related C_3_ and C_4_ species under fluctuating light, first the steady state light response of photosynthetic gas exchange was measured under 21% and 2% O_2_ to minimize photorespiration (**Figure 1**). We focus here on photosynthetic parameters at the light intensities that were used in subsequent fluctuating light treatments, 800 and 100 µmol m^-2^ s^-1^ PFD (**Table 1**). Two-way ANOVA was used to assess the effects of photosynthetic pathway, oxygen concentration, and their interactions within each genus (**Table 2**). At both light intensities C_3_ species trended towards higher assimilation values and quantum yields under photorespiration- suppressing (2% O_2_) conditions than C_4_ species.

**Figure 1:**
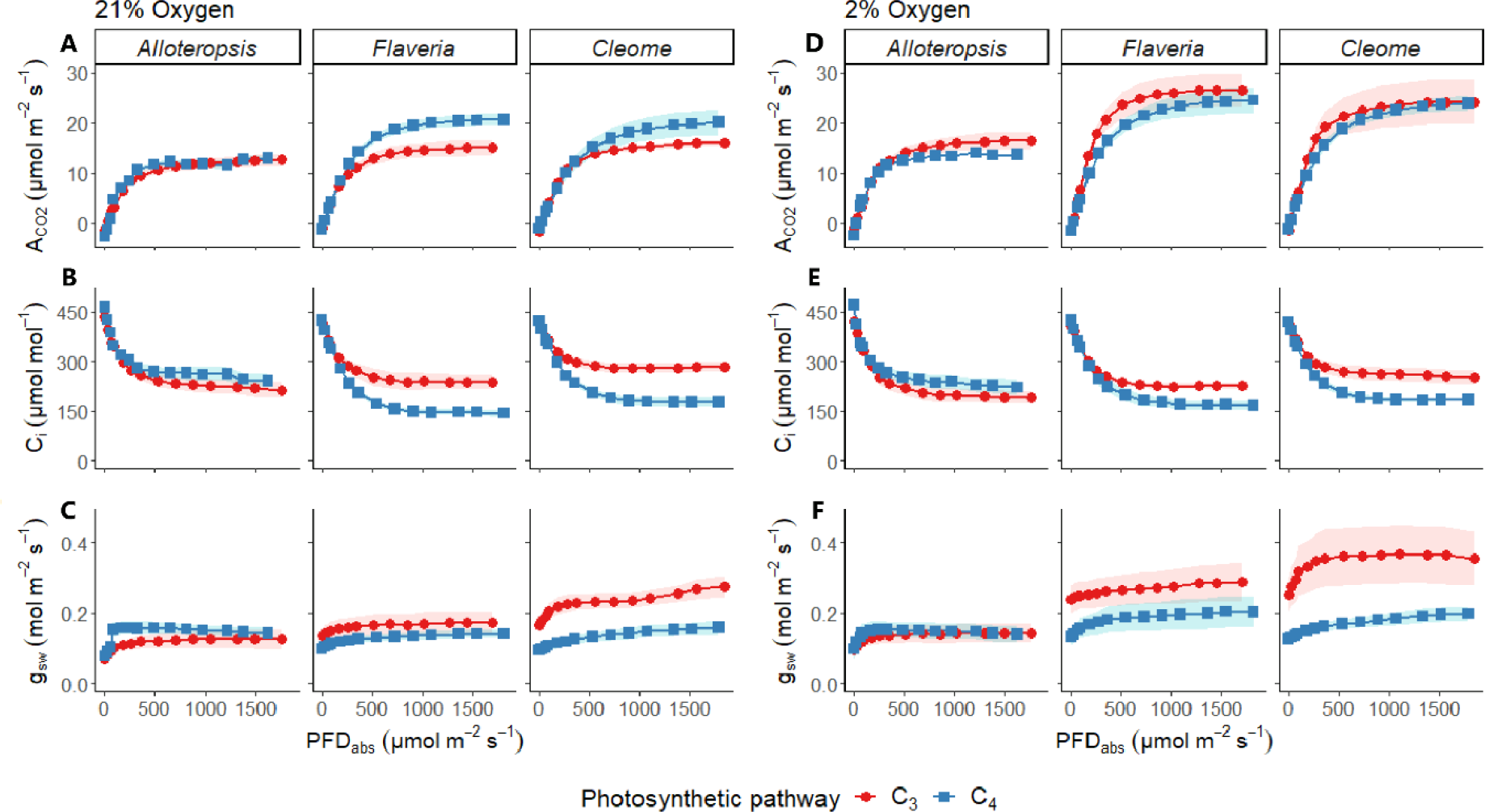
Light response curves of phylogenetically linked C_3_ and C_4_ *Alloteropsis*, *Flaveria* and *Cleome* species at 21% and 2% O_2_. As a function of absorbed light intensity, plots show net CO_2_ assimilation (A_CO2_, **A, D**), intercellular CO_2_ concentration (C_i_, **B, E**), and stomatal conductance to water vapour (g_sw_, **C, F**). Ribbons represent standard error of the mean (n=5).

**Table 1:** Photosynthetic parameters estimated from steady state light response curves under 21% and 2% O_2_ on phylogenetically linked C_3_ and C_4_ *Alloteropsis, Flaveria* and *Cleome* species. Respiration in the light *(Rd)* was calculated by fitting the light response curves with a non-rectangular hyperbola. Values for *800* A_CO2_, *100* A_CO2_, C_i_ *800*, C_i_ *100, 800* ΦCO_2_, and *100* ΦCO_2_ were taken at PFD = 800 µmol m^-2^ s^-1^ and PFD = 100 µmol m^-2^ s^-1^. Means and standard error of the mean are shown (n = 5).

| Genus | Parameter | 21% Oxygen |  | 2% Oxygen |  |
| --- | --- | --- | --- | --- | --- |
|  |  | C<sub>3</sub> | C<sub>4</sub> | C<sub>3</sub> | C<sub>4</sub> |
| *Alloteropsis* | 800 A<sub>CO2</sub> (μmol m<sup>-2</sup> s<sup>-1</sup>) | 11.3 ± 0.9 | 12.3 ± 0.5 | 15.0 ± 1.2 | 13.1 ± 0.1 |
|  | 100 A<sub>CO2</sub> (μmol m<sup>-2</sup> s<sup>-1</sup>) | 3.2 ± 0.5 | 4.7 ± 0.5 | 4.8 ± 0.7 | 4.7 ± 0.7 |
|  | C<sub>i800</sub> (μmol mol<sup>-1</sup>) | 235 ± 20 | 265 ± 15 | 207 ± 21 | 245 ± 20 |
|  | C<sub>i100</sub> (μmol mol<sup>-1</sup>) | 348 ± 9 | 349 ± 2 | 334 ± 9 | 345 ± 2 |
|  | 800 ΦCO<sub>2</sub> | 0.017 ± 0.003 | 0.021 ± 0.002 | 0.025 ± 0.006 | 0.022 ± 0.00 |
|  | 100 ΦCO<sub>2</sub> | 0.047 ± 0.012 | 0.075 ± 0.014 | 0.065 ± 0.005 | 0.076 ± 0.018 |
|  | *Rd* | 0.6 ± 0.1 | 1.7 ± 0.3 | 1.5 ± 0.3 | 1.5 ± 0.1 |
| *Flaveria* | 800 A<sub>CO2</sub> (μmol m<sup>-2</sup> s<sup>-1</sup>) | 13.8 ± 1.4 | 18.7 ± 1.0 | 25.0 ± 2.9 | 21.5 ± 1.7 |
|  | 100 A<sub>CO2</sub> (μmol m<sup>-2</sup> s<sup>-1</sup>) | 3.9 ± 0.2 | 4.2 ± 0.3 | 6.6 ± 0.3 | 4.7 ± 0.4 |
|  | C<sub>i800</sub> (μmol mol<sup>-1</sup>) | 246 ± 25 | 157 ± 8 | 230 ± 8 | 182 ± 18 |
|  | C<sub>i100</sub> (μmol mol<sup>-1</sup>) | 352 ± 6 | 339 ± 4 | 351 ± 5 | 343 ± 9 |
|  | 800 ΦCO<sub>2</sub> | 0.022 ± 0.004 | 0.027 ± 0.00 | 0.038 ± 0.01 | 0.030 ± 0.005 |
|  | 100 ΦCO<sub>2</sub> | 0.057 ± 0.005 | 0.057 ± 0.007 | 0.085 ± 0.01 | 0.056 ± 0.009 |
|  | *Rd* | 0.4 ± 0.4 | 0.8 ± 0.3 | 0.6 ± 0.2 | 0.4 ± 0.2 |
| *Cleome* | 800 A<sub>CO2</sub> (μmol m<sup>-2</sup> s<sup>-1</sup>) | 14.6 ± 0.5 | 23.5 ± 2.0 | 27.3 ± 4.0 | 27.4 ± 0.9 |
|  | 100 A<sub>CO2</sub> (μmol m<sup>-2</sup> s<sup>-1</sup>) | 4.1 ± 0.2 | 3.2 ± 0.4 | 6.2 ± 0.6 | 4.7 ± 0.5 |
|  | C<sub>i800</sub> (μmol mol<sup>-1</sup>) | 281 ± 14 | 190 ± 12 | 266 ± 21 | 192 ± 9 |
|  | C<sub>i100</sub> (μmol mol<sup>-1</sup>) | 363 ± 7 | 352 ± 3 | 358 ± 18 | 344 ± 7 |
|  | 800 ΦCO<sub>2</sub> | 0.020 ± 0.001 | 0.026 ± 0.006 | 0.033 ± 0.012 | 0.030 ± 0.003 |
|  | 100 ΦCO<sub>2</sub> | 0.048 ± 0.004 | 0.056 ± 0.011 | 0.090 ± 0.016 | 0.061 ± 0.012 |
|  | *Rd* | 1.5 ± 1.0 | 1.2 ± 0.4 | 2.1 ± 0.7 | 0.8 ± 0.5 |

**Table 2:** ANOVA table of light response curve parameters at 100 and 800 µmol m^-2^ s^-1^ PFD for phylogenetically linked C_3_ and C_4_ *Alloteropsis, Flaveria* and *Cleome* species. Photosynthetic pathway, PP. O_2_ concentration, [O_2_]. Interaction effect, PP:[O_2_]. Table shows degrees of freedom; *F*-value; and *p*-value. Significant *p*-values (*a* < 0.05) are shown in bold.

|  | *Alloteropsis* |  |  | *Flaveria* |  |  | *Cleome* |  |  |
| --- | --- | --- | --- | --- | --- | --- | --- | --- | --- |
| Parameter | PP | $[\text{O}_2]$ | PP:[$\text{O}_2$] | PP | $[\text{O}_2]$ | PP:[$\text{O}_2$] | PP | $[\text{O}_2]$ | PP:[$\text{O}_2$] |
| $800 A_{\text{CO}_2}$ | 1,16;<br>0.35;<br>0.56 | 1,16;<br>7.77;<br>**0.01** | 1,16;<br>3.24;<br>0.09 | 1,16;<br>0.12;<br>0.73 | 1,16;<br>13.48;<br>**0.01** | 1,16;<br>4.83;<br>**0.04** | 1,15;<br>0.01;<br>0.91 | 1,15;<br>6.18;<br>**0.02** | 1,15;<br>0.82;<br>0.28 |
| $100 A_{\text{CO}_2}$ | 1,16;<br>1.21;<br>0.29 | 1,16;<br>1.60;<br>0.22 | 1,16;<br>1.56;<br>0.23 | 1,16;<br>7.01;<br>**0.02** | 1,16;<br>25.64;<br>**$\leq 0.001$** | 1,16;<br>11.38;<br>**0.01** | 1,15;<br>7.28;<br>**0.02** | 1,15;<br>15.06;<br>**0.01** | 1,15;<br>0.43;<br>0.52 |
| $C_{i\ 800}$ | 1,16;<br>3.21;<br>0.09 | 1,16;<br>1.62;<br>0.22 | 1,16;<br>0.05;<br>0.83 | 1,16;<br>17.63;<br>**$\leq 0.001$** | 1,16;<br>0.09;<br>0.76 | 1,16;<br>1.63;<br>0.22 | 1,16;<br>31.44;<br>**$\leq 0.001$** | 1,16;<br>0.19;<br>0.66 | 1,16;<br>1.33;<br>0.58 |
| $C_{i\ 100}$ | 1,16;<br>0.87;<br>0.36 | 1,16;<br>1.96;<br>0.40 | 1,16;<br>0.74;<br>0.40 | 1,16;<br>2.52;<br>0.13 | 1,16;<br>0.11;<br>0.75 | 1,16;<br>0.16;<br>0.70 | 1,16;<br>3.65;<br>0.07 | 1,16;<br>1.01;<br>0.33 | 1,16;<br>0.04;<br>0.84 |
| $800 \Phi \text{CO}_2$ | 1,16;<br>0.22;<br>0.07 | 1,16;<br>8.12;<br>**0.01** | 1,16;<br>4.47;<br>**0.05** | 1,16;<br>0.13;<br>0.72 | 1,16;<br>11.90;<br>**0.01** | 1,16;<br>5.95;<br>**0.03** | 1,16;<br>0.19;<br>0.67 | 1,16;<br>6.96;<br>**0.02** | 1,16;<br>2.18;<br>0.15 |
| $100 \Phi \text{CO}_2$ | 1,16;<br>11.03;<br>**0.01** | 1,16;<br>2.67;<br>0.12 | 1,16;<br>2.04;<br>0.17 | 1,16;<br>15.56;<br>**0.01** | 1,16;<br>13.93;<br>**0.01** | 1,16;<br>16.53;<br>**$\leq 0.001$** | 1,16;<br>4.50;<br>**0.05** | 1,16;<br>21.01;<br>**$\leq 0.001$** | 1,16;<br>12.72;<br>**0.01** |

At 800 µmol m^-2^ s^-1^ PFD, net CO_2_ assimilation (A_CO2_) values in C_4_ *A. semialata MDG* (12.3 ± 0.5 µmol m^-2^ s^-1^), C_4_ *F. bidentis* (18.7 ± 1.0 µmol m^-2^ s^-1^), and C_4_ *G. gynandra* (23.5 ± 2.0 µmol m^-2^ s^-1^) were respectively higher than phylogenetic pairs C_3_ *A. semialata GMT* (11.3 ± 0.9 µmol m^-2^ s^-1^), C_3_ *F. cronquistii* (13.8 ± 1.4 µmol m^-2^ s^-1^), and C_3_ *T. hassleriana* (14.6 ± 0.5 µmol m^-2^ s^-1^). Under 2% oxygen, photosynthetic rates increased by 33% in C_3_ *A. semialata GMT*, 81% in C_3_ *F. cronquistii,* and 86% in C_3_ *T. hassleriana*, compared to a more modest increase of 6%, 15% and 16% in their respective C_4_ pairs, leading to significant effects of O_2_ at 800 µmol m^-2^ s^-1^ PFD on A_CO2_ as well as on quantum yield of CO_2_ assimilation (ΦCO_2_) across all three genera (*p* ≤ 0.02).

At 100 µmol m^-2^ s^-1^ PFD, the effects of photosynthetic pathway, O_2_ concentration and their interaction on A_CO2_ varied between genera. In *Alloteropsis,* none of the effects were significant, although C_4_ *A. semialata MDG* (4.7 ± 0.5 µmol m^-2^ s^-1^) showed slightly higher A_CO2_ values than C_3_ *A. semialata GMT* (3.2 ± 0.5 µmol m^-2^ s^-1^) under ambient O_2_. In *Flaveria,* values of A_CO2_ were similar under ambient O_2_ in C_4_ *F. bidentis* (4.2 ± 0.3 µmol m^-2^ s^-1^), and C_3_ *F. cronquistii* (3.9 ± 0.2 µmol m^-2^ s^-1^), but significantly higher in C_3_ *F. cronquistii* under 2% O_2_ (4.7 ± 0.4 vs 6.6 ± 0.3 µmol m^-2^ s^-1^, respectively). As a result, both the main effects of O_2_ and photosynthetic pathway as well as their interaction on both A_CO2_ and ΦCO_2_ were significant in *Flaveria* (*p* ≤ 0.02). In *Cleome*, A_CO2_ under 21% O_2_ was lower in C_4_ *G. gynandra* (3.2 ± 0.4 µmol m^-2^ s^-1^) than in C_3_ *T. hassleriana* (4.1 ± 0.2 µmol m^-2^ s^-1^) and both rates increased under 2% O_2_ (4.7 ± 0.5 and 6.2 ± 0.6 µmol m^-2^ s^-1^, respectively), resulting in significant effects of both photosynthetic pathway and O_2_ concentration (*p* ≤ 0.02).

### CO_2_ assimilation under fluctuating light differs between genera and is significantly affected by fluctuation frequency, photosynthetic pathway, and oxygen concentration

After characterising steady state light responses, CO_2_ assimilation rates were measured in response to three different 1-hour fluctuating light treatments. Each treatment consisted of acclimation at 150 µmol m^-2^ s^-1^ PFD, followed by repetitive stepwise changes between 800 and 100 µmol m^-2^ s^-1^ PFD where each light step lasted 6, 30, or 300 seconds. Data from minutes 50-60 of each treatment (**Figure 2**) were used for analysis to exclude the effect of initial induction (for the complete timeseries see **Supplementary figure 1**). The fluctuating light response generally consisted of a rise in A_CO2_ towards steady state in the 800 µmol m^-2^ s^-1^ PFD period, and a subsequent decrease during the 100 µmol m^-2^ s^-1^ PFD period. A_CO2_ was strongly increased under 2% O_2_ in the C_3_ species, but much less so in the C_4_ species. More subtle patterns varied by photosynthetic type, fluctuation length, and between genera as described below.

**Figure 2:**
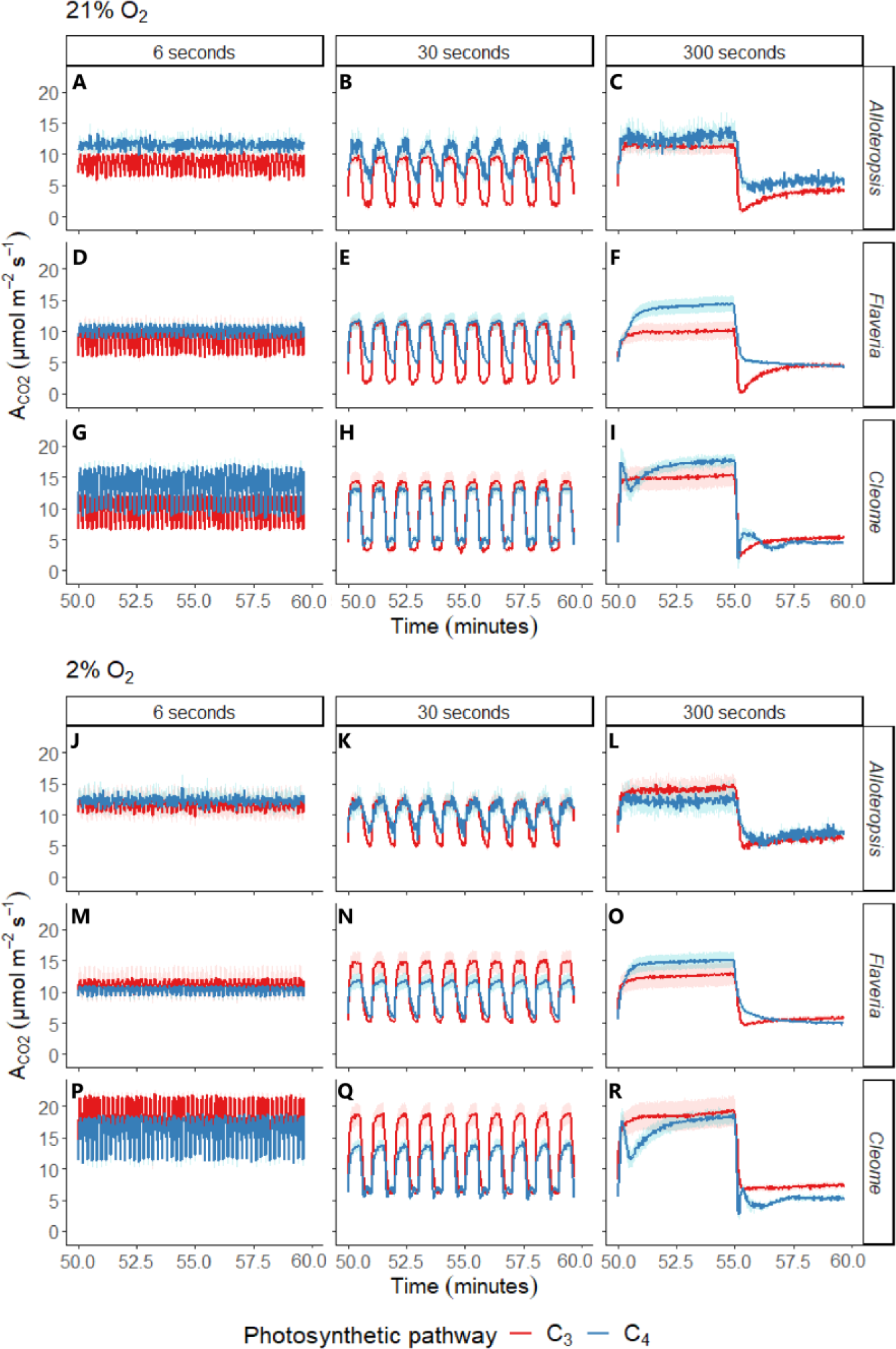
Net CO_2_ assimilation (A_CO2_) in phylogenetically linked C_3_ and C_4_ *Alloteropsis*, *Flaveria* and *Cleome* species under three different fluctuating light regimes at 21% and 2% O_2_. Each light regime consisted of alternating 800 and 100 µmol m^-2^ s^-1^ PFD periods, where each light step lasted 6, 30, or 300 seconds before changing. Treatments were started after leaves were acclimated at 150 µmol m^-2^ s^-1^ PFD and lasted 1 hour, data was analysed from minutes 50-60 of each treatment. Ribbons represent standard error of the mean (n=5). The full timeseries is shown in **Supplementary** figure 1.

After transitioning from 800 to 100 µmol m^-2^ s^-1^ PFD, A_CO2_ in C_4_ *A. semialata MDG* and C_4_ *F. bidentis* decreased more gradually compared to the immediate drop followed by a gradual rise towards steady state observed in C_3_ *A. semialata GMT,* C_3_ *F. cronquistii* and C_3_ *T. hassleriana*. This instantaneous drop in assimilation in the C_3_ species at the start of the 100 µmol m^-2^ s^-1^ PFD period is commonly attributed to photorespiration (Forrester et al., 1966, Wynn et al., 1973), and indeed was suppressed in all C_3_ species under 2% oxygen (most easily seen in **Figure 2 C/F/I** compared to **Figure 2 L/O/R)**. The slower decrease of A_CO2_ in C_4_ *A. semialata MDG* and C_4_ *F. bidentis* was evident under 30 s (**Figure 2 B/E**) and 300 s light steps (**Figure 2 C/F**) in both oxygen concentrations, whilst under 6 s light steps (**Figure 2 A/D**) A_CO2_ at 100 µmol m^-2^ s^-1^ stayed closer to rates obtained during the 800 µmol m^-2^ s^-1^ PFD periods, suggesting a less significant initial decline. Irrespective of oxygen concentration and unlike the delayed decrease observed in the other C_4_ species, in C_4_ *G. gynandra* an initial dip in A_CO2_ was observed immediately following the transition to low light (**Figure 2 I/R**).

Following the transition from 100 to 800 µmol m^-2^ s^-1^ PFD, induction patterns strongly varied between the three C_4_ species, in contrast with the more consistent patterns observed in C_3_ species. C_4_ *F. bidentis* and C_4_ *G. gynandra* had higher A_CO2_ during the 300 s light steps than C_3_ *F. bidentis* and C_3_ *T. hassleriana* (**Figure 2 C/F**), but similar or lower A_CO2_ than their C_3_ counterparts during the 30 s light step (**Figure 2 B/E**) even under 21% oxygen, suggesting a comparatively greater lag in photosynthetic induction. However, induction of A_CO2_ in C_4_ *A. semialata MDG* was very similar to that of C_3_ *A. semialata GMT*. In C_4_ *G. gynandra* a strong temporary depression in A_CO2_ was observed after an initial sharp increase upon exposure to higher light under the 300 s light step that was not affected by the suppression of photorespiration by low oxygen (see **Figure 2 I vs R**).

To compare photosynthetic performance between C_3_ and C_4_ species whilst accounting for their different steady state photosynthetic capacities, A_CO2_ was expressed as a relative percentage of steady state values obtained from light response curves (percentages across a single fluctuation are shown in **Figure 3**. For the corresponding carbon assimilation values see **Supplementary figure 2**, which were compared to the steady state values in **Table 2/Figure 1**). This analysis showed clear, systematic differences between C_3_ and C_4_ species during the 100 µmol m^-2^ s^-1^ PFD steps, where all C_4_ species were able to sustain higher A_CO2_ under 100 µmol m^-2^ s^-1^ relative to steady state than their matching C_3_ counterparts under both 21% and 2% oxygen. However, no clear trend between photosynthetic types was apparent during the 800 µmol m^-2^ s^-1^ PFD steps.

**Figure 3:**
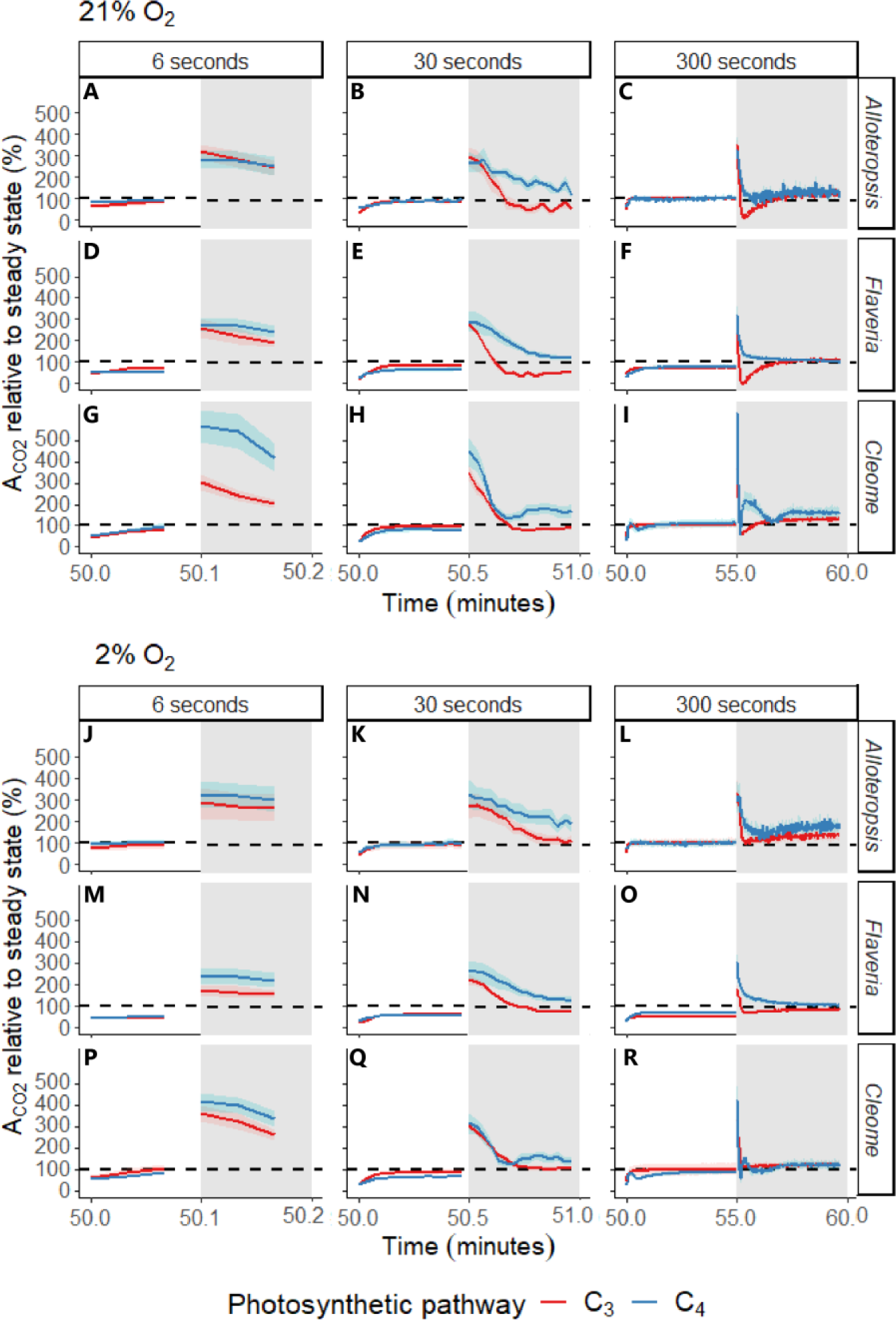
Net CO_2_ assimilation (A_CO2_) relative to steady state (%) across the 800 and 100 µmol m^-2^ s^-1^ PFD light steps of differing length starting at the 50 minute mark, in white and grey respectively. Depending on the fluctuating light treatment, subplots are showing one complete fluctuation of 12, 60, or 600 seconds. Values represent A_CO2_ at a given point in the fluctuating light treatment relative to A_CO2_ obtained from steady state light response curves at the light intensity of each period in phylogenetically linked C_3_ and C_4_ *Alloteropsis*, *Flaveria* and *Cleome* species at 21% and 2% O_2_. The dashed line represents 100%, where assimilation would be exactly that of steady state. Ribbons represent standard error of the mean (n=5). The corresponding absolute assimilation values are provided in **Supplementary** figure 2.

### Stimulation of CO_2_ assimilation at low light phase of fluctuations is most prominent in short light steps and significantly greater in C_4_ compared to C_3_ species

To quantify the stimulation of A_CO2_ during the 100 µmol m^-2^ s^-1^ PFD steps of the fluctuations, the average A_CO2_ during the 100 µmol m^-2^ s^-1^ PFD steps was normalized against the steady state rate at the same intensity (**Figure 4 D-F**, for a boxplot of the absolute A_CO2_ values see **Supplementary figure 3 E-G**). This analysis showed that assimilation was higher than steady state (as seen in the **Figure 3** grey half, values greater than 100%) in all species immediately following the transition but declined with duration of the light steps (**Table 3**, *p* ≤ 0.001 for all). In addition, the relative stimulation compared to steady state values was consistently significantly greater in the C_4_ species compared to the C_3_ species in the *Flaveria* and *Cleome* pairs (*Flaveria p* ≤ 0.01; *Cleome p* ≤ 0.001). Although not significant, a similar trend was observed for the *Alloteropsis* pair at 300 s and 30 s, but not at 6 s light steps.

**Figure 4:**
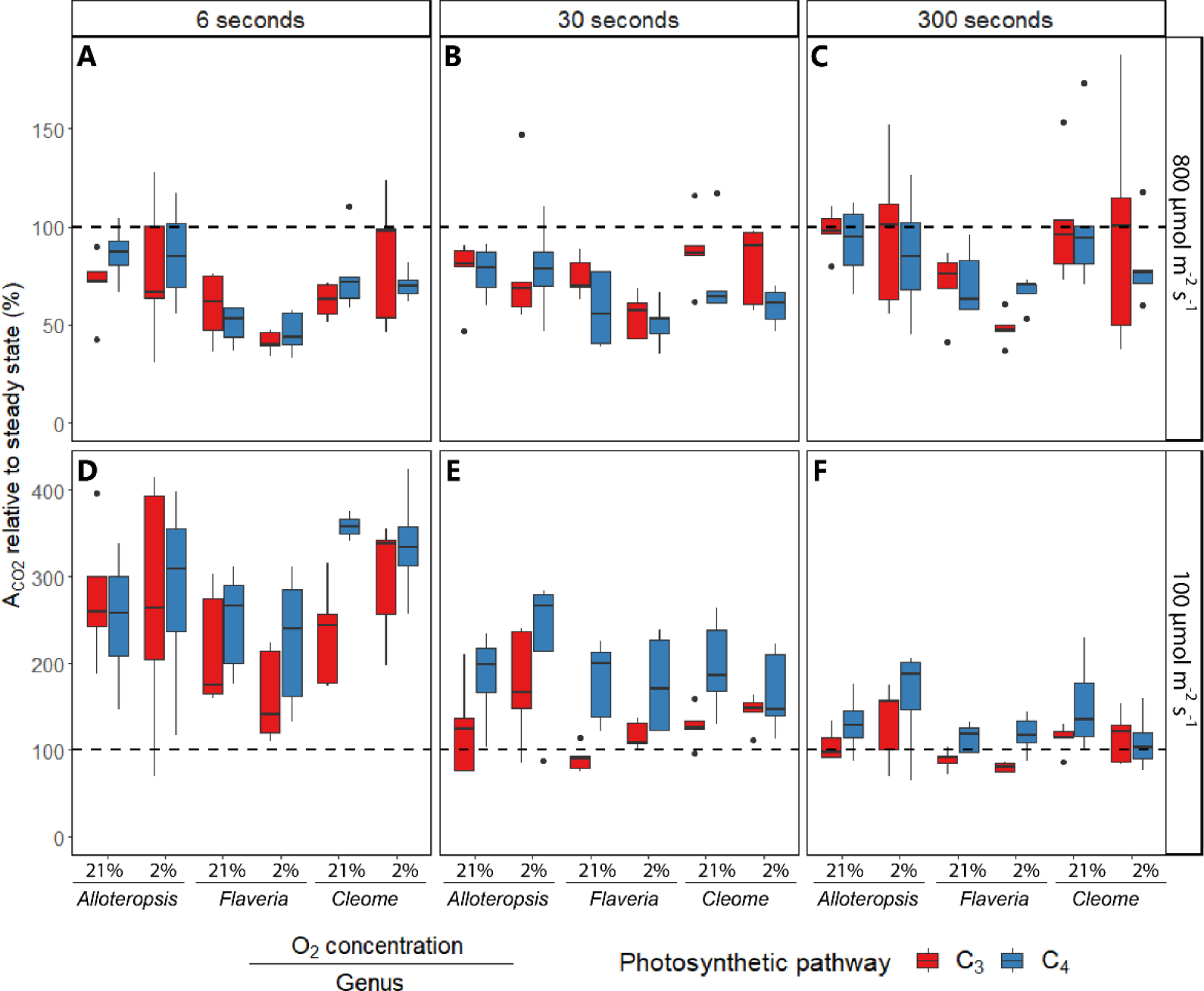
Boxplots of net CO_2_ assimilation (A_CO2_) relative to steady state (%) under the 800 and 100 µmol m^-2^ s^-1^ PFD light steps of the fluctuating light regimes. Each regime consisted of alternating 800 and 100 µmol m^-2^ s^-1^ PFD periods, where each light step lasted 6, 30, or 300 seconds. For each period, A_CO2_ across the timeseries for phylogenetically linked C_3_ and C_4_ *Alloteropsis*, *Flaveria* and *Cleome* species at 21% or 2% O_2_ was calculated as a percentage of steady state values obtained from light response curves at the same light intensity and O_2_ concentration. The dashed line represents 100%, where assimilation rate would equal steady state. Box edges represent the lower and upper quartiles, the solid line indicates the median, and points represent outliers beyond 1.5 times the interquartile range (n = 5 for each combination of species/oxygen). Three-way ANOVA (**Table 4**) was used to test the effect of photosynthetic pathway, fluctuating length, O_2_ concentration and their interaction on A_CO2_ relative to steady state in *Alloteropsis, Flaveria*, and *Cleome*. The corresponding absolute assimilation values are shown in **Supplementary** figure 3.

**Table 3:**
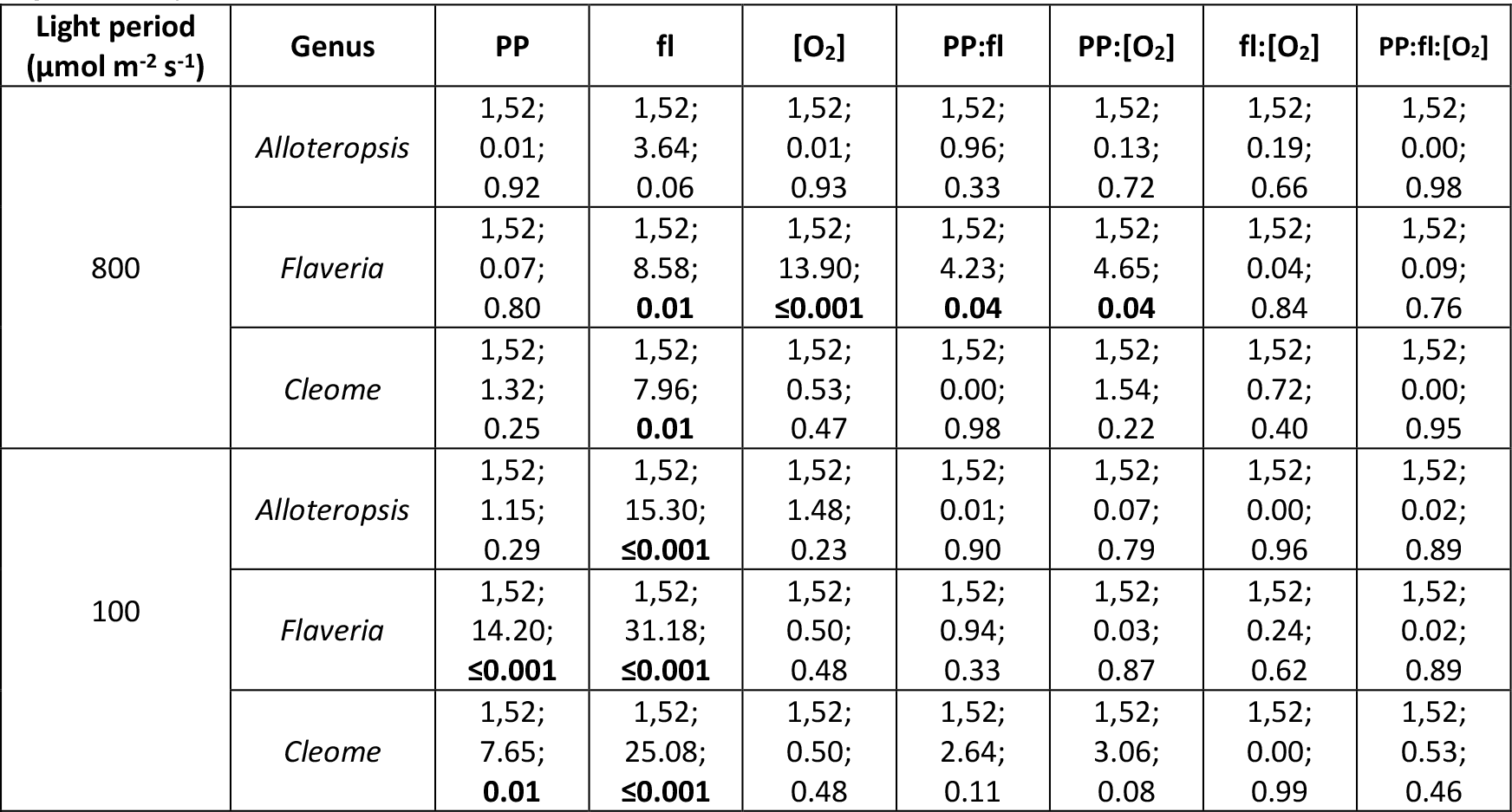
ANOVA table of percentage A_CO2_ relative to steady state during the two different light steps of the light fluctuation treatments for phylogenetically linked C_3_ and C_4_ *Alloteropsis*, *Flaveria* and *Cleome* species. Photosynthetic pathway, PP. Fluctuating length, fl. O_2_ concentration, [O_2_]. Interaction effects, PP:fl, PP:[O_2_], fl:[O_2_], and PP:fl:[O_2_]. Table shows degrees of freedom, *F*-value, and *p*-value. Significant *p*-values (*a* < 0.05) are shown in bold.

**Table 4:** ANOVA table of ΦCO_2_ during the two different light steps of the light fluctuation treatments for phylogenetically linked C_3_ and C_4_ *Alloteropsis, Flaveria* and *Cleome* species. Photosynthetic pathway, PP. Fluctuating length, fl. O_2_ concentration, [O_2_]. Interaction effects, PP:fl, PP:[O_2_], fl:[O_2_], and PP:fl:[O_2_]. Table shows degrees of freedom, *F*-value, and *p*-value. Significant *p*-values (*a* < 0.05) are shown in bold.

| Light period<br>($\mu\text{mol m}^{-2} \text{s}^{-1}$) | Genus | PP | fl | $[\text{O}_2]$ | PP:fl | PP:$[\text{O}_2]$ | fl:$[\text{O}_2]$ | PP:fl:$[\text{O}_2]$ |
| --- | --- | --- | --- | --- | --- | --- | --- | --- |
| 800 | *Alloteropsis* | 1,52;<br>0.00;<br>0.97 | 1,52;<br>2.00;<br>0.16 | 1,52;<br>7.08;<br>**0.01** | 1,52;<br>0.76;<br>0.39 | 1,52;<br>1.22;<br>0.27 | 1,52;<br>0.20;<br>0.66 | 1,52;<br>0.00;<br>0.96 |
|  | *Flaveria* | 1,52;<br>0.10;<br>0.75 | 1,52;<br>2.67;<br>0.11 | 1,52;<br>0.02;<br>0.89 | 1,52;<br>1.17;<br>0.28 | 1,52;<br>4.56;<br>**0.03** | 1,52;<br>0.40;<br>0.53 | 1,52;<br>0.77;<br>0.38 |
|  | *Cleome* | 1,52;<br>0.01;<br>0.92 | 1,52;<br>11.15;<br>**0.01** | 1,52;<br>16.33;<br>**$\leq 0.001$** | 1,52;<br>0.53;<br>0.47 | 1,52;<br>4.45;<br>**0.04** | 1,52;<br>2.48;<br>0.12 | 1,52;<br>0.17;<br>0.68 |
| 100 | *Alloteropsis* | 1,52;<br>5.70;<br>**0.02** | 1,52;<br>23.37;<br>**$\leq 0.001$** | 1,52;<br>14.02;<br>**$\leq 0.001$** | 1,52;<br>0.25;<br>0.62 | 1,52;<br>0.94;<br>0.34 | 1,52;<br>0.04;<br>0.85 | 1,52;<br>0.07;<br>0.79 |
|  | *Flaveria* | 1,52;<br>0.01;<br>0.94 | 1,52;<br>34.60;<br>**$\leq 0.001$** | 1,52;<br>1.28;<br>0.26 | 1,52;<br>0.00;<br>0.99 | 1,52;<br>3.26;<br>0.08 | 1,52;<br>1.15;<br>0.70 | 1,52;<br>0.17;<br>0.68 |
|  | *Cleome* | 1,52;<br>1.04;<br>0.31 | 1,52;<br>38.70;<br>**$\leq 0.001$** | 1,52;<br>9.40;<br>**0.02** | 1,52;<br>1.21;<br>**0.04** | 1,52;<br>1.81;<br>0.28 | 1,52;<br>1.75;<br>0.19 | 1,52;<br>0.42;<br>0.52 |

Three-way ANOVA (**Table 3**) showed A_CO2_ relative to steady state was significantly affected by fluctuation length in all genera (*p* ≤ 0.001 for all), as well as by photosynthetic pathway in *Flaveria* and *Cleome* (*Flaveria p* ≤ 0.01; *Cleome p* ≤ 0.001). Overall, although all species had greater CO_2_ fixation during the 100 µmol m^-2^ s^-1^ PFD periods than under steady state, the effect was time sensitive and therefore more significant during shorter light steps. In addition, C_4_ species were able to sustain the higher rates for longer than their C_3_ counterparts. The greatest increases in relative assimilation occurred during the 6 s light steps (**Figure 4 D**) where C_4_ species were on average 329% of steady state A_CO2_ compared to 242% in C_3_ species under 21% oxygen and similarly 290% in C_4_ vs 243% in C_3_ at 2% oxygen. Under 30 s light steps (**Figure 4 E**) the stimulation of A_CO2_ relative to steady state was less pronounced than 6 s, but still significantly higher in C_4_ species at 187% compared to C_3_ species at 114% of steady state A_CO2_ at 21% O_2_, and 190% vs 146% of steady state A_CO2_ at 2% O_2_, respectively. The impact of C_4_ photosynthesis was most apparent under these two light steps, whereas during the 300 s light steps (**Figure 4 F**) the stimulation of A_CO2_ was generally less evident. Averaged across the 300 s, C_4_ species were operating at 132% relative to steady state A_CO2_ compared to C_3_ species at 103% under ambient oxygen, and at 130% and 109% of steady state A_CO2_ under 2% O_2_, respectively. Interestingly, A_CO2_ relative to steady state across all three light step durations was typically higher in C_4_ compared to C_3_ species under both 21% and 2% oxygen and no significant effect of the interaction between photosynthetic pathway and oxygen concentration was found in any of the genera, suggesting a systematic advantage to C_4_ photosynthesis to bridge low light periods which was not explained by suppression of photorespiration.

To estimate to what extent the low light stimulation of CO_2_ assimilation was decoupled from photochemical provision of ATP and NADPH, ΦCO_2_ was calculated for each light step (**Figure 5, Table 4**). Based on steady state stoichiometry of electron flow and proton requirements for ATP synthesis and NADPH:ATP energy demands, the theoretical maximum ФCO_2_ has been estimated as 0.111 CO_2_/photon for C_3_ species (Ehleringer and Pearcy, 1983), as 0.064 CO_2_/photon for C_4_ NADP-ME and NAD-ME species accounting for BS leakiness (Yin and Struik, 2018), and due to the theorised lower energy requirements of mixed C_4_ pathways as 0.075 CO_2_/photon for mixed subtype NADP-ME-PEPCK (Ishikawa et al., 2016, Yin and Struik, 2021). Here we consider observations of quantum yields exceeding these theoretical maxima as conservative estimates for the degree of decoupling. At 6 s light steps, C_3_ *F. cronquistii* and C_3_ *A. semialata GMT* stayed well below the theoretical maximum, but the ФCO_2_ values of C_3_ *T. hassleriana* were significantly higher. ФCO_2_ values of C_4_ *A. semialata MDG,* C_4_ *F. bidentis, and* C_4_ *G. gynandra* during the lower light periods of the 6 s and 30 s fluctuations were also significantly higher than the theoretical limit, suggesting that the provision of ATP and reductant was not directly coupled to production from the thylakoid reactions. By comparing the ФCO_2_ values with these theoretical maxima, it is possible to estimate the degree of decoupling in units of fixed CO_2_/photon. At 6 s light steps, the theoretical limit was exceeded by 0.043 ± 0.019 CO_2_/photon in C_4_ *A. semialata MDG*, 0.022 ± 0.008 CO_2_/photon in C_4_ *F. bidentis,* 0.091 ± 0.008 CO_2_/photon in C_4_ *G. gynandra* and 0.042 ± 0.008 CO_2_/photon in C_3_ *T. hassleriana*. At 30 s light steps, these numbers decreased to 0.021 ± 0.011 CO_2_/photon in C_4_ *A. semialata MDG*; 0.008 ± 0.005 CO_2_/photon in C_4_ *F. bidentis*; and 0.005 ± 0.003 CO_2_/photon in C_4_ *G. gynandra*.

**Figure 5:**
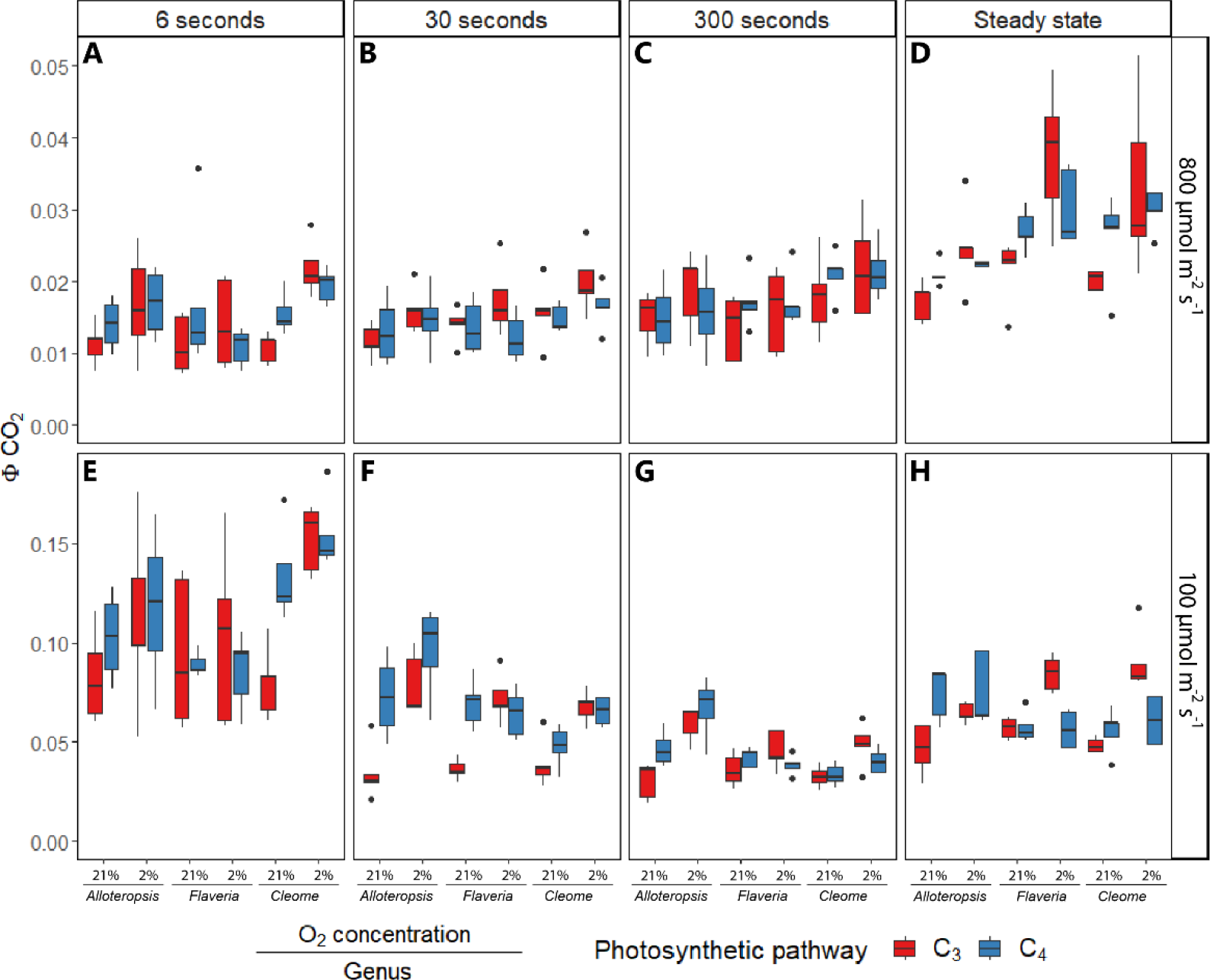
Boxplots of the quantum yield of carbon assimilation (ΦCO_2_) under the 800 and 100 µmol m^-2^ s^-1^ PFD periods of the fluctuating light regimes or from steady state measurements. Each fluctuating regime consisted of alternating 800 and 100 µmol m^-2^ s^-1^ PFD periods, where each light step lasted 6, 30, or 300 seconds. Box edges represent the lower and upper quartiles, the solid line indicates the median, and points represent outliers beyond 1.5 times the interquartile range (n = 5 for each combination of species/measurement condition). Three-way ANOVA was used to test the effect of photosynthetic pathway, fluctuating length, O_2_ concentration and their interaction on ΦCO_2_ in *Alloteropsis, Flaveria*, and *Cleome* (results shown in **Table 5**).

**Table 5:** Phylogenetically linked C_3_ and C_4_ *Alloteropsis*, *Flaveria*, and *Cleome* species used in this study with their photosynthetic pathway, class, and main C_4_ subtype if applicable.

| Genus | Species and accession | Photosynthetic pathway | C4 subtype | Class |
| --- | --- | --- | --- | --- |
| *Alloteropsis* | *Alloteropsis semialata* subspecies *semialata* accession GMT | C<sub>3</sub> | --- | Monocot |
|  | *Alloteropsis semialata* subspecies *eckloniana* accession MDG | C<sub>4</sub> | NADP-ME-PEPCK |  |
| *Flaveria* | *Flaveria cronquistii* | C<sub>3</sub> | --- | Dicot |
|  | *Flaveria bidentis* | C<sub>4</sub> | NADP-ME |  |
| *Cleome* | *Tarenaya hassleriana* | C<sub>3</sub> | --- |  |
|  | *Gynandropsis gynandra* | C<sub>4</sub> | NAD-ME |  |

### Depression of CO_2_ assimilation at high light phase of fluctuations is not significantly affected by photosynthetic pathway

Unlike in the 100 µmol m^-2^ s^-1^ PFD periods there was no clear trend between the C_3_ and C_4_ response at 800 µmol m^-2^ s^-1^ PFD periods (**Figure 4 A-C**, for a boxplot of the corresponding A_CO2_ values see **Supplementary figure 3 A-C**).

A_CO2_ values during the 800 µmol m^-2^ s^-1^ PFD light steps were lower than under steady state (below 100% line in **Figure 3**). Three-way ANOVA (**Table 3**) was used to analyse the effects of light step duration, photosynthetic pathway, oxygen concentration, and their interactions on relative A_CO2_. None of these were significant for the *Alloteropsis* subspecies. In *Flaveria*, weakly significant interactions between light step duration and photosynthetic pathway (*p* = 0.04), as well as between oxygen concentration and photosynthetic pathway (*p* = 0.04) were observed, indicating a more complex conditional impact of photosynthetic pathway on assimilation rate relative to steady state. In C_4_ *F. bidentis* relative A_CO2_ gradually increased with light step duration, whereas in C_3_ *F. cronquistii* this increase was only observed between 6 and 30 s but had reached a plateau between 30 and 300 s. Whilst relative A_CO2_ was depressed by 2% oxygen in both *Flaveria* species, the effect was more pronounced in C_3_ *F. cronquistii*. In *Cleome*, light step duration significantly impacted relative A_CO2_ (*p* = 0.01) which increased with duration in both C_3_ and C_4_ species.

Quantum yields during the 800 µmol m^-2^ s^-1^ PFD periods across all fluctuation regimes and oxygen concentrations were lower than steady state across all species (**Figure 5 A-D**), indicating reduced efficiency of carbon assimilation. In *Alloteropsis*, oxygen concentration significantly impacted ФCO_2_ (*p* = 0.01, **Table 4**), with 2% oxygen being associated with higher values. In *Flaveria* and *Cleome* 2% oxygen was associated with higher quantum yields only in the C_3_ species, and lower or similar values in their C_4_ counterparts (significant interactions between O_2_ and photosynthetic pathway *p* ≤ 0.05).

## Discussion

### C_4_ species are better able to sustain photosynthetic rates than C_3_ species during the low light phase of fluctuations

The effect of fluctuating light on C_4_ relative to C_3_ photosynthesis was systematically evaluated in three phylogenetically controlled comparisons using repetitive low and high light steps with three contrasting durations. The results support the hypothesis that C_4_ species are better able to sustain photosynthetic rates than C_3_ species during the lower light periods of fluctuating light. The theoretical basis for this hypothesis suggests that the large metabolite pools necessary for diffusional transfer between M and BS in C_4_ photosynthesis and the reversible reactions linking these metabolic intermediates can work as a capacitor, providing greater flexibility to respond to variations in light intensity (Leegood and von Caemmerer, 1989, Stitt and Zhu, 2014). In this study, although all tested species had generally higher carbon assimilation during the 100 µmol m^-2^ s^-1^ PFD periods relative to steady state, C_4_ *A. semialata MDG,* C_4_ *F. bidentis*, and C_4_ *G. gynandra* were elevated higher above steady state than C_3_ *A. semialata GMT*, C_3_ *F. bidentis,* and C_3_ *T. hassleriana* under the same fluctuating light regime (**Figure 4 D-F**). As C_4_ species had higher relative assimilation under both 21% and 2% oxygen, the greater stimulation under low light cannot be solely attributed to increased presence of photorespiration in C_3_ species during fluctuating light, and the prevalence of this result across species from diverse evolutionary lineages and C_4_ subtypes suggests that this ability to sustain high photosynthetic rates after a transition to lower light may indeed be linked to other general features of the C_4_ pathway – such as the large metabolite pools intrinsic to CCM operation (Arrivault et al., 2017, Leegood and Furbank, 1984, Lilley et al., 1977, Stitt et al., 1985).

Metabolic pools also play a role in C_3_ species, and post-illumination CO_2_ fixation has previously been attributed to pools of C_3_ intermediates, and ATP and redox equivalents that accumulate during higher light fluctuations (Kaiser et al., 2015). In C_3_ *A. semialata GMT*, C_3_ *F. bidentis,* and C_3_ *T. hassleriana*, the higher A_CO2_ values relative to steady state observed during 6 s fluctuations, as well as during the first half of the 30 s fluctuations (**Figure 3 BEH & KNQ**) are in line with previous observations of carbon assimilation exceeding steady state rates immediately after sunflecks (Pons and Pearcy, 1992, Sharkey et al., 1986). However, metabolic pools in C_3_ species are typically considerably smaller than in C_4_ species (Borghi et al., 2022), which may be why C_4_ *A. semialata MDG,* C_4_ *F. bidentis*, and C_4_ *G. gynandra* were able to sustain higher relative rates for longer than their C_3_ counterparts (**Figure 3**), as evidenced by the higher average rates across longer fluctuations (**Figure 4 D-F**). However, the capacity of C_4_ species to buffer through transitions to lower light still decreases across time as metabolite pools are depleted (Slattery et al., 2018), and carbon assimilation relative to steady state during those periods was inversely related to the length of the fluctuations. Altogether, the higher and more sustained stimulation of C_4_ photosynthetic rates compared to C_3_ rates at low light is consistent with prior work (Laisk and Edwards, 1997, Lee et al., 2022, Li et al., 2021) and explains why studies using different fluctuating light regimes can yield contrasting estimates for the comparative advantage of C_4_ versus C_3_ photosynthesis.

Finally, the quantum yield of photosynthesis provides another indication of the storage capacity of C_4_ metabolic pools. In C_4_ species ФCO_2_ during the 6 s and 30 s fluctuations was consistently above the theoretical maximum (Ehleringer and Pearcy, 1983, Ishikawa et al., 2016, Morales et al., 2018, Yin and Struik, 2018, Yin and Struik, 2021), indicating that the energetic equivalent of fixing ∼0.022-0.091 CO_2_/photon under 6 s fluctuations and ∼0.005-0.021 CO_2_/photon under 30 s fluctuations was being supplied outside of the thylakoid light reactions. These numbers provide a conservative estimate of the extent of decoupled CO_2_ fixation in C_4_ species, which may have been sustained by redox equivalents from malate decarboxylation, with demands for ATP and NADH being buffered through reversible reactions linking 3-PGA and TP, or interconversion of 3-PGA and PEP (Arrivault et al., 2017, Bräutigam et al., 2008, Slattery et al., 2018, Stitt and Zhu, 2014, Wang et al., 2014b). Leaf-level and canopy simulations emphasise ФCO_2_ as the largest determinant of photosynthesis in the lower canopy (Bellasio and Farquhar, 2019, Gu et al., 2014) and the stimulation of low light ФCO_2_ as observed here at three contrasting fluctuation frequencies could provide an important mitigation mechanism of the intrinsically lower maximum quantum yield of C_4_ plants under low light (Medeiros et al., 2022, Ubierna et al., 2011).

### The C_4_ response during the transition to higher light is related to the specific subtype metabolism

The comparative high light performance of C_4_ photosynthesis was not uniform across the three genera, and instead was impacted by the specific characteristics of the C_4_ pathway within each genus. Light fluctuations have been suggested to hinder the formation of the large metabolite pools necessary for optimal CCM operation, resulting in impaired suppression of photorespiration and lagging photosynthetic induction during metabolite build-up (Sage and McKown, 2006). Indeed, we previously found that the C_4_ species analysed here were all slower to induce photosynthesis from darkness relative to their C_3_ counterparts (Arce Cubas et al., 2023). However, theoretical work indicates that not all subtypes are equally reliant on gradients: mixed C_4_ pathways like NADP-ME-PEPCK do not need metabolite gradients as large as NADP-ME or NAD-ME subtypes, as mixed subtypes can concurrently use different transfer acids (Wang et al., 2014a). The use of both Mal and Asp shuttles also allows for finer regulation of the ATP:NADPH ratio in response to changes in light, as only Mal transport brings redox equivalents into the BS (Yin and Struik, 2021). Consistent with this interpretation, the highest A_CO2_ relative to steady state rates in C_4_ species during the 800 µmol m^-2^ s^-1^ PFD periods were found in C_4_ *A. semialata MDG* which is suggested to rely on a mixed NADP-ME-PEPCK C_4_ pathway (Ueno and Sentoku, 2006) (**Figure 4 A-C**). The lack of effect of light step duration on A_CO2_ during the high light periods in *Alloteropsis* (**Table 3**) can be explained by photosynthetic induction during these fluctuating light regimes being relatively fast (**Figure 2 ABC**). In contrast, high light A_CO2_ relative to steady state was significantly lower with shorter fluctuations in both *Cleome* species; whereas in *Flaveria,* C_4_ *F. bidentis* relative assimilation was more significantly reduced during shorter fluctuations than in C_3_ *F. cronquistii*, indicating the C_4_ cycle lagged behind C_3_ activation.

The genus-specific rather than photosynthetic pathway effects detailed above in combination with the varying impact of light step duration provides an important consideration for interpretation of previous work. Li et al. (2021) compared a selection of six C_4_ species with eight C_3_ species, concluding that C_4_ species utilized fluctuating light less efficiently. Our results suggest the slow induction observed in that study could be associated with the selected C_4_ species being mostly of the NADP-ME subtype (five out of six, one of which was *F. bidentis*) and one NAD-ME subtype. Indeed, Lee et al. (2022) recently reported distinct species-specific induction patterns across six C_4_ grass species. Interestingly, Lee et al. concluded that their selection of C_4_ species assimilated more carbon under fluctuating light in comparison with six C_3_ species. Aside from the inclusion of different species in both papers, these seemingly opposite conclusions could also be due to a slight difference in fluctuating light treatments. Although both experiments had low light steps of two minutes, high light steps were two minutes in the Li et al. (2021) study and four minutes in Lee et al. (2022). Since both studies observed a slower decrease in photosynthetic rates in C_4_ species after the transition to lower light, the additional two minutes of higher light in the Lee et al. study may have reduced the comparative penalty of C_4_ induction relative to the benefits of higher assimilation during the lower light periods, explaining the contrasting conclusions.

### Fluctuations in light cause CO_2_ bursts in C_4_ G. gynandra

C_4_ *G. gynandra* had distinctive assimilation kinetics after each light transition in the 300 s fluctuations (**Figure 2 I&R**). At the start of the 800 µmol m^-2^ s^-1^ PFD period, a rapid increase in A_CO2_, described in previous work as a CO_2_ gulp (Laisk and Edwards, 1997), was immediately followed by a CO_2_ burst, with another CO_2_ burst upon changing to the 100 µmol m^-2^ s^-1^ PFD period. These bursts were not a product of photorespiration, as unlike the CO_2_ bursts observed after a transition to lower light in C_3_ species, in C_4_ *G. gynandra* they occurred independent of oxygen concentration.

Previous studies on sunflecks (Laisk and Edwards, 1997) have characterised the post-illumination CO_2_ burst as a specific feature of the NAD-ME and PEPCK pathways, and other fluctuating light studies have also reported it primarily for NAD-ME species (Lee et al., 2022). Unlike NADP-ME species, where malate decarboxylation is linked to reducing equivalents from the C_3_ cycle, in NAD-ME species the C_3_ and C_4_ cycles are less tightly coupled – oxaloacetate is first reduced to malate and then decarboxylated in the mitochondria, but the redox balance is uncoupled from the C_3_ cycle (Ishikawa et al., 2016). This can result in excess CO_2_ being released despite insufficient RuBP regeneration upon a transition to lower light, causing unfixed CO_2_ to leak out of the BS and A_CO2_ to drop.

The CO_2_ gulp and burst at the start of the 800 µmol m^-2^ s^-1^ PFD period is similar to induction kinetics observed in short dark-light fluctuations in NAD-ME *Amaranthus cruentus* (Laisk and Edwards, 1997). These were attributed to formation of alanine from the decarboxylation of aspartate in low light (or darkness), leading to rapid conversion of alanine to pyruvate followed by phosphorylation to PEP when light is increased (Laisk and Edwards, 1997, Lee et al., 2022). Based on this interpretation. the initial PEP carboxylation following the increase in light would exceed the rate at which PEP pools can be replenished, but subsequently crashes and readjusts while PEP regeneration is re-established. This may account for the observed temporary gulp and subsequent steady increase in A_CO2_. Furthermore, the centripetal chloroplast positioning found in NAD-ME species (Yoshimura et al., 2004) could increase path length for metabolites and CO_2_, and lead to a more pronounced form of the biphasic induction previously observed in NADP-ME species (Lee et al., 2022).

## Conclusions

The presented work compared C_4_ to C_3_ photosynthesis in response to fluctuating light. By using three independent phylogenetically controlled comparisons and fluctuations with three contrasting light step durations the presented work circumvented issues in previous studies to yield more robust conclusions. The results showed that the stimulation of A_CO2_ in the low light phase was both higher and more sustained in C_4_ photosynthesis across all three comparisons, suggesting this could be a common comparative advantage of C_4_ photosynthesis. In contrast, observed patterns of A_CO2_ in the high light phase were found to be largely genus-specific, rather than attributable to photosynthetic pathway.

## Materials and methods

### Plant materials

To control for evolutionary distance, three pairs of phylogenetically linked *Alloteropsis*, *Flaveria*, and *Cleome* C_3_ and C_4_ species (shown in **Table 5**, see also Arce Cubas et al., 2023) were selected. Significant evolutionary distance exists between the selected genera and C_4_ photosynthesis evolved independently in each. The C_4_ origin dates back approximately 17 million years (Ma) in *Cleome*, ∼ 2 Ma in *Flaveria*, and is even more recent in *Alloteropsis* (Christin et al., 2011, Lundgren et al., 2015). In addition, the selected species include both monocots (*Alloteropsis*) and dicots (*Flaveria* and *Cleome*), and all three major decarboxylase enzymes of the C_4_ pathway: NADP-ME-PEPCK (C_4_ *A. semialata MDG*), NAD-ME (C_4_ *F. bidentis*), and NAD-ME (C_4_ *G. gynandra*) (Bräutigam et al., 2008, Gowik et al., 2011, Ueno and Sentoku, 2006).

### Plant growth and propagation

*Flaveria* and *Cleome* species were grown in a Conviron walk-in growth room (Conviron Ltd., Winnipeg, MB, CA) at 20 °C, 60% relative humidity (RH), and 150 µmol m^-2^ s^-1^ PFD over a 16-hour photoperiod; and the *Alloteropsis* accessions in a glasshouse in Cambridge, England, at 18-25 °C, 40-60% RH, with supplemental lightning to provide a minimum of 140-160 µmol m^-2^ s^-1^ PFD in addition to incoming irradiance. All plants were well-watered and grown in Levington Advance M3 compost (Scotts, Ipswich, UK) mixed with Miracle-Gro All Purpose Continuous Release Osmocote (Scotts Miracle-Gro Company, Marysville, OH, USA; 4 L compost : 25 g Osmocote), with vermiculite being added to the *Alloteropsis* soil mix to prevent waterlogging (1 L vermiculite : 4 L compost : 25 g Osmocote).

*Alloteropsis* and *Flaveria* species were vegetatively propagated whilst *Cleome* species were grown from seed. *Alloteropsis* MDG and GMT accession tillers were grown in 2 L pots and all gas exchange measurements taken after 2 weeks. For *Flaveria* propagation, lateral shoot cuttings were dipped in Doff Hormone Rooting Powder (Doff Portland Ltd., Hucknall, UK) to induce root development, grown on 0.25 L pots, and measured after 8-10 weeks. *F. cronquistii* requires vegetative propagation, so *F. bidentis* plants were first grown from seed and subsequently propagated via cuttings*. Cleome* germination was induced with a 30 °C/20 °C day/night cycle for *T. hassleriana*, and at 30 °C for *G. gynandra*. The germinated seeds were sown in 24-cell trays before transfer to 0.25 L pots. Due to the different developmental rates of the *Cleome* species, germination was staggered so both species could be measured at approximately the same developmental stage, after 8-10 weeks for *G. gynandra* and 4-6 weeks for *T. hassleriana*. All plants were measured during vegetative state.

### Gas exchange measurements at 21% and 2% O_2_

Gas exchange under steady and fluctuating light conditions was measured on young, fully expanded leaves using an open gas exchange system (LI-6800, LI-COR, Lincoln, NE, USA) with a Multiphase Flash Fluorometer (MPF) chamber (6800-01A, LI-COR). Chamber conditions were controlled at 410 ppm sample CO_2_ concentration, 60% relative humidity with average leaf VPD of 1.3 ± 0.1 kPa, 25 °C heat exchanger temperature, and flow rate of 600 µmol s^-1^. Actinic light was provided by the MPF and composed of 90% red (625 nm) and 10% blue light (475 nm).

For experiments in 2% O_2_, a pre-mixed 2% O_2_ and 98% N2 gas mixture (BOC Ltd., Woking, UK) was supplied to the LI-6800 through the air inlet using a mass flow controller (EL-FLOW, Bronkhorst Hight-tech BV, Ruurlo, NL) and an open T-junction to regulate constant surplus flow according to manufacturer instructions. The LI-6800 Infrared Gas Analyser (IRGA) calibration was adjusted to the O_2_ concentration in the instrument constants prior to measurement.

### Steady state light response curves

Photosynthetic responses to steady light were measured for all species at both 21% and 2% O_2_. Leaves were illuminated with 1000 µmol m^-2^ s^-1^ PFD for 20-40 minutes to allow CO_2_ assimilation and stomatal conductance to reach steady state, and gas exchange was subsequently measured in a descending gradient of light intensity: 2000, 1700, 1500, 1200,

1000, 800, 600, 400, 300, 200, 100, 75, 30 and 0 µmol m^-2^ s^-1^ PFD. Gas exchange parameters were logged between 120 – 240 seconds at a given light intensity, when leaf intracellular CO_2_ concentration (C_i_) and CO_2_ assimilation were stable.

Respiration in the light (*Rd*) was estimated from the y-intercept of a linear regression of the slope before the inflection point. Measurements at 0 and 30 µmol m^-2^ s^-1^ PFD were not included in the regression to account for the Kok effect (Kok, 1949). Photosynthetic rates and C_i_ at 100 and 800 µmol m^-2^ s^-1^ PFD were taken as the steady state values for comparison with fluctuating light measurements. ΦCO_2_ was calculated using **Equation 1**.

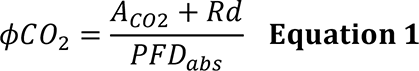

### Fluctuating light experiments and analysis

To measure photosynthetic responses to fluctuating light, leaves were first acclimated at 150 µmol m^-2^ s^-1^ PFD for 30-60 minutes until stomatal conductance and photosynthetic rates reached constant levels. Using a custom program, leaves were then exposed to repetitive stepwise fluctuations in light intensity from 800 to 100 µmol m^-2^ s^-1^ PFD for 1 hour, with gas exchange parameters recorded every 2 seconds. Three different light treatments were tested, with each light step lasting 6, 30, or 300 seconds. To avoid interference with the shorter fluctuations and the data sampling interval the averaging time of the LI-6800 was set to zero, as was additional averaging time, and the IRGAs were only matched prior to the program starting. All fluctuating light treatments were conducted at both 21% and 2% O_2_.

As measurements during light fluctuations violate the steady state assumption underlying default rate equations, a storage flux term for CO_2_ and H2O was computed from changes in cuvette concentration between timepoints and applied to adjust carbon assimilation and transpiration rates – and consequently to other instrument calculations derived from those values – using **Equation 2 and 3**.

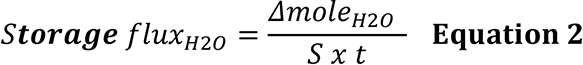

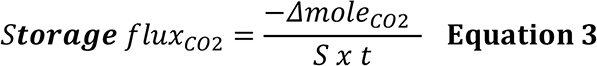

The Δ change in moles of CO_2_ or H2O was calculated from the cuvette volume 8.67e^-5^ m^3^ using the ideal gas law with instrument recordings of current and previous gas concentration, pressure, and air temperature, and converted to storage flux with leaf area (S) and time since last log (t). The signum reconciles CO_2_ flux with the consensus of uptake flux being positive.

### Leaf absorptance

After gas exchange measurements, the spectral qualities of the leaves were measured with an integrating sphere (LI-1800-12, LI-COR) optically connected to a miniature spectrometer (STS-VIS, Ocean Insight, Orlando, FL, USA) following manufacturer instructions (LI-COR, 1988). Leaf absorptance (Labs) was calculated using **Equation 4**, where Ts and Rs are transmittance and reflectance of a diffuse sample.

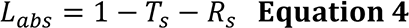

For the light response curves, incident PFD was converted to absorbed PFD (PFD_abs_) using Labs of the red and blue emission wavelengths of the 6800-01A MPF light source. For the specific absorptance values, see **Supplementary table 1**.

### Data processing

Data from the last 10 minutes of the fluctuating light treatment were used for further analysis. To analyse the relative performance of each species, net photosynthesis (A_CO2_) under fluctuating light was expressed as a percentage of the steady state rates achieved at the corresponding light intensity. Additionally, ΦCO_2_ under fluctuating light was calculated using **Equation 1**.

The area under the curve (AUC) (Makowski et al., 2019) of A_CO2_ relative to steady state and ΦCO_2_ during the 100 and 800 µmol m^-2^ s^-1^ PFD periods of the light treatment was integrated using the trapezoidal rule (Jawień, 2014) and divided by the duration to obtain an average value for each light level which was used to compare between fluctuations of different length and, in the case of ΦCO_2_, directly to steady state values.

### Statistical analysis

Each phylogenetically controlled comparison was run as an independent experiment, thus statistical analyses were conducted separately on paired *Alloteropsis*, *Flaveria,* and *Cleome* light response curves (*800* A_CO2_, *100* A_CO2_, C_i_ 800, C_i_ 100, *800* ΦCO_2_, *100* ΦCO_2_, *Rd*), and fluctuating light measurements (A_CO2_ relative to steady state, ΦCO_2_). Two-way ANOVA was used to test for the effects of photosynthetic pathway, oxygen concentration, and their interactions on steady state photosynthesis parameters; and three-way ANOVA to test for the effects of photosynthetic pathway, fluctuation length, oxygen concentration, and their interactions on A_CO2_ relative to steady state and ΦCO_2_. Specifically for the ΦCO_2_ analysis, the quantum yields obtained under steady state at 800 and 100 µmol m^-2^ s^-1^ PFD were included in the dataset as an additional fluctuation length. For each ANOVA, assumptions of normality, homogeneity of variance and sphericity were satisfied. Mean and standard error of the mean for steady state photosynthesis parameters, A_CO2_ relative to steady state, and ΦCO_2_ across the light fluctuation regimes were calculated for reporting.

All data analysis and plot generation was done with R 4.1.1 (R Core Team, 2021) on RStudio 2022.12.0+353 (Posit Team, 2022) using the tidyverse (Wickham et al., 2019), RColorBrewer (Neuwirth, 2014), lme4 (Bates et al., 2015), and bayestestR libraries (Makowski et al., 2019).

## Acknowledgements

Lucía Arce Cubas was jointly funded by The Cambridge Commonwealth, European & International Trust; and by Mexico’s Consejo Nacional de Ciencia y Tecnología (CONACyT). This work was supported by the BBSRC via grant BB/T007583/1 awarded to Dr. Johannes Kromdijk. The authors also thank Dr. Pascal-Antoine Christin at The University of Sheffield, Dr. Marjorie Lundgren at Lancaster Environmental Centre, and Prof. Peter Westhoff at Heinrich Heine University Düsseldorf for respectively providing the original *A. semialata GMT* and *MDG* accessions, *F. cronquistii* cuttings, and *F. bidentis* seeds. For the purpose of open access, the authors have applied a Creative Commons Attribution (CC BY) licence to any Author Accepted Manuscript version arising from this submission.

## Author contributions

JK and LAC conceived the study. JK, LAC and ACB designed the experiments. LAC carried out all experiments, data analysis and interpretation, and drafted the manuscript. CRGS helped with the fluctuating light experimental setup and data interpretation. RLV helped with the 2% O_2_ experimental setup and provided support with gas exchange experiments. ELB procured the initial plant material. All authors contributed to the article and approved the submitted version.

