## Supplemental Fig 1-3 and Table 1 for "Lessons from relatives: C4 photosynthesis enhances CO_2_ assimilation during the low-light phase of fluctuations"

Supplementary material

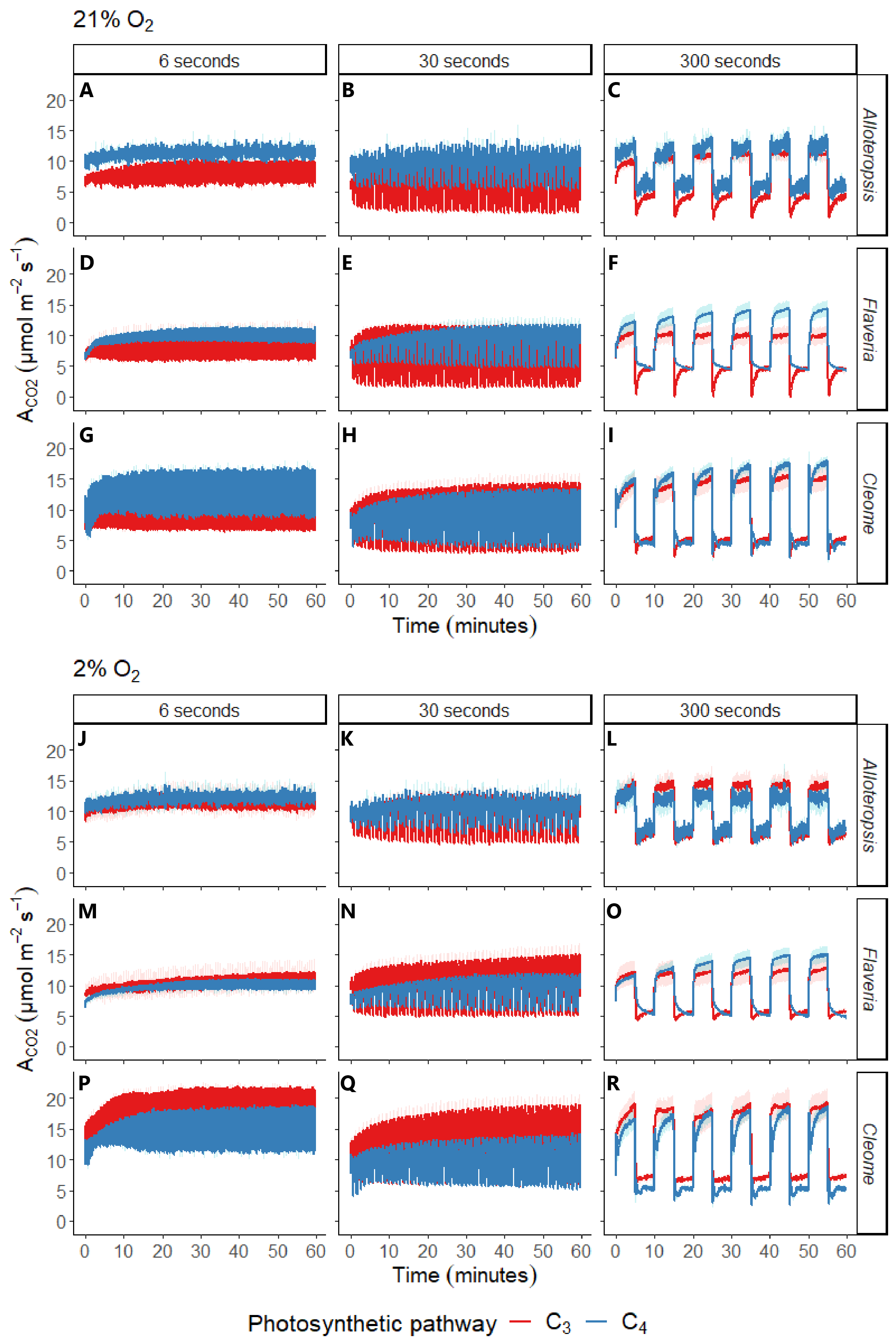

**Supplementary figure 1:** Net CO_2_ assimilation (A_CO2_) in phylogenetically linked C_3_ and C_4_ *Alloteropsis*, *Flaveria* and *Cleome* species under three different fluctuating light regimes at 21% and 2% O_2_. Each light regime consisted of alternating 800 and 100 µmol m^-2^ s^-1^ PFD periods, where each light step lasted 6, 30, or 300 seconds before changing. Treatments were started after leaves were acclimated at 150 µmol m^-2^ s^-1^ PFD and lasted 1 hour, but data in the study was taken from minutes 50-60 of each treatment. Ribbons represent standard error of the mean (n=5).

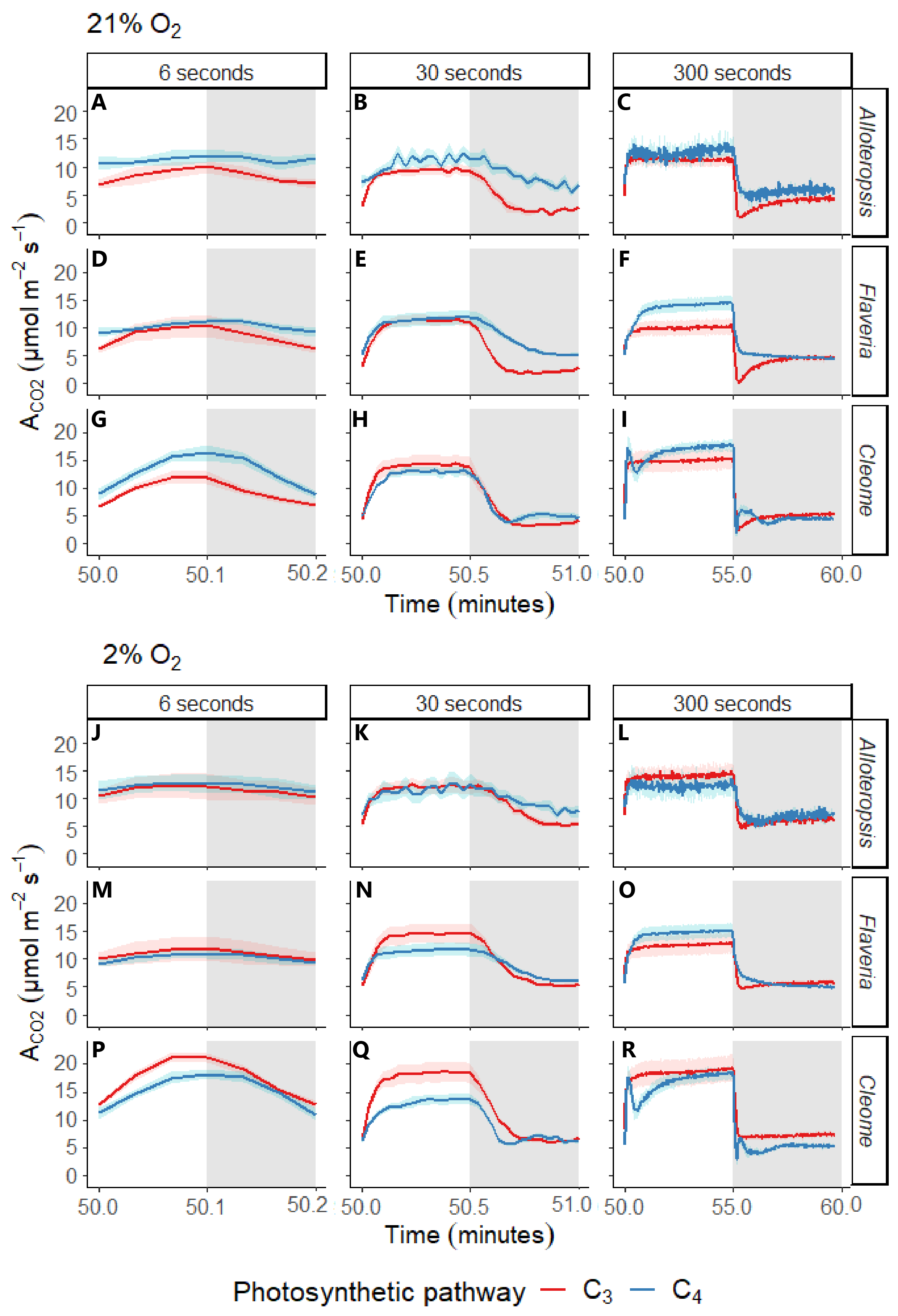

**Supplementary figure 2:** Net CO_2_ assimilation (A_CO2_) across the 800 and 100 µmol m^-2^ s^-1^ PFD light steps, in white and grey respectively. All data was taken starting from 50 min into the experiment. Depending on the fluctuating light treatment, each light step was 6, 30, or 300 seconds. Values represent phylogenetically linked C_3_ and C_4_ *Alloteropsis*, *Flaveria* and *Cleome* species at 21% and 2% O_2_. Ribbons represent standard error of the mean (n=5). The corresponding A_CO2_ relative to steady state (%) values are in **Figure 4**.

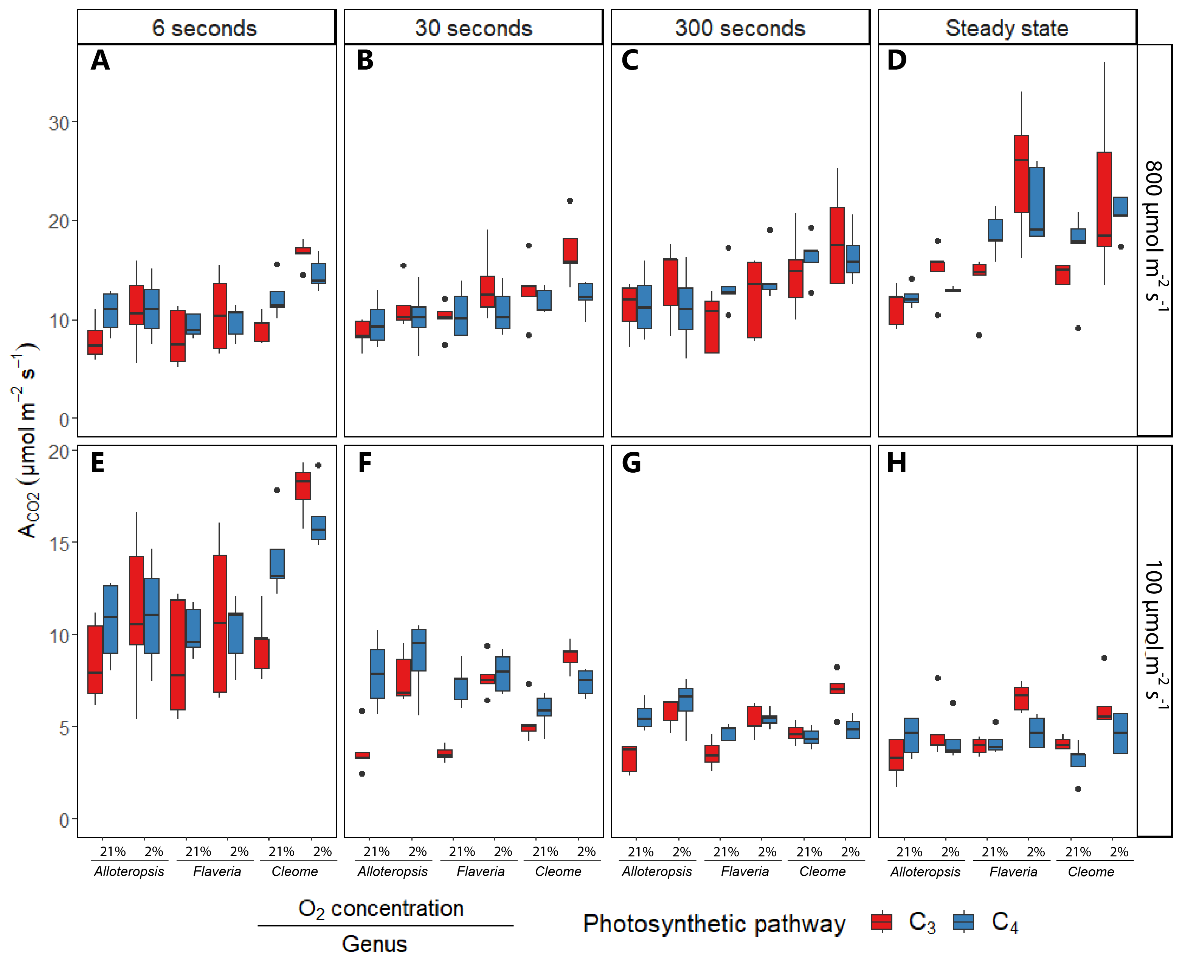

**Supplementary figure 3:** Boxplots of net carbon assimilation (A_CO2_) under the 800 and 100 µmol m^-2^ s^-1^ PFD periods of the fluctuating light regimes. Each regime consisted of alternating 800 and 100 µmol m^-2^ s^-1^ PFD periods, where each light step lasted 6, 30, or 300 seconds before changing. The plot includes A_CO2_ steady state values taken from the light response curves. The area under the curve (AUC) A_CO2_ of for each period across the timeseries for phylogenetically linked C_3_ and C_4_ *Alloteropsis*, *Flaveria* and *Cleome* species at 21% or 2% O_2_ was calculated from between minutes 50-60 of each fluctuating light treatment and converted to a rate for ease of comparison. Box edges represent the lower and upper quartiles, the solid line indicates the median, and points represent outliers beyond 1.5 times the interquartile range (n = 5 for each combination of species/measurement condition).

| **Supplementary table 1:** Leaf absorptance values of phylogenetically linked C_3_ and C_4_ *Alloteropsis*, *Flaveria*, and *Cleome* species from the blue (475 nm) and red (625 nm) wavelengths of the actinic light source used in experiments, measured with an integrating sphere. Means and standard error of the mean are shown (n = 5). | | | |
| --- | --- | --- | --- |
| **Genus** | **Species** | **L_abs_ (475 nm)** | **L_abs_ (625 nm)** |
| *Alloteropsis* | C3 *A. semialata GMT* | 0.90±0.02 | 0.88±0.02 |
|  | C4 *A. semialata MDG* | 0.87±0.02 | 0.81±0.02 |
| *Flaveria* | C3 *F. cronquistii* | 0.88±0.02 | 0.85±0.02 |
|  | C4 *F. bidentis* | 0.94±0.00 | 0.90±0.00 |
| *Cleome* | C3 *T. hassleriana* | 0.95±0.00 | 0.92±0.00 |
|  | C4 *G. gynandra* | 0.93±0.00 | 0.89±0.00 |
